# No unexpected CRISPR-Cas9 off-target activity revealed by trio sequencing of gene-edited mice

**DOI:** 10.1101/263129

**Authors:** Vivek Iyer, Katharina Boroviak, Mark Thomas, Brendan Doe, Edward Ryder, David Adams

## Introduction

CRISPR-Cas technologies have transformed genome-editing of experimental organisms and have immense therapeutic potential. Despite significant advances in our understanding of the CRISPR-Cas9 system, concerns remain over the potential for off-target effects. Recent studies have addressed these concerns using whole-genome sequencing (WGS) of gene-edited embryos or animals to search for *de novo* mutations (DNMs), which may represent candidate changes induced by poor editing fidelity[1–3]. Critically, these studies used strain-matched but not pedigree-matched controls and thus were unable to reliably distinguish generational or colony-related differences from true DNMs. Here we used a trio design and whole genome sequenced 8 parents and 19 embryos, where 10 of the embryos were mutagenised with well-characterised gRNAs targeting the coat colour Tyrosinase *(Tyr)* locus.

## Results and Discussion

To perform our analysis, we chose two gRNAs targeting exon 2 of *Tyr*, Tyr2F and Tyr2R, that had typical off-target scores similar to that of the sgRNA used by Schaefer et al. [3](Supplementary Methods). *Tyr* is responsible for black coat colour and eye pigmentation in C57BL/6 mice[4], so its disruption should not be detrimental to embryonic development. The CRISPR-treated group was split to include five embryos treated with Tyr2F and five embryos treated with Tyr2R, while three untreated embryos from each of the three control groups (“Cas9 only”, “No injection” and “Sham injection”) were also collected (**Fig. 1a**). Microinjections were performed into the cytoplasm of 1-cell zygotes[5], which were then briefly cultured to assess viability and then transferred into 0.5 day post coital (d.p.c) pseudopregnant females. Embryos in the “Sham injection” group were microinjected with water only, and the “Cas9 only” embryos were microinjected with Cas9 protein only. All embryos were harvested at 12.5 d.p.c (Supplementary Table 1) and genomic DNA from both parents and embryos extracted.

**Figure 1:**
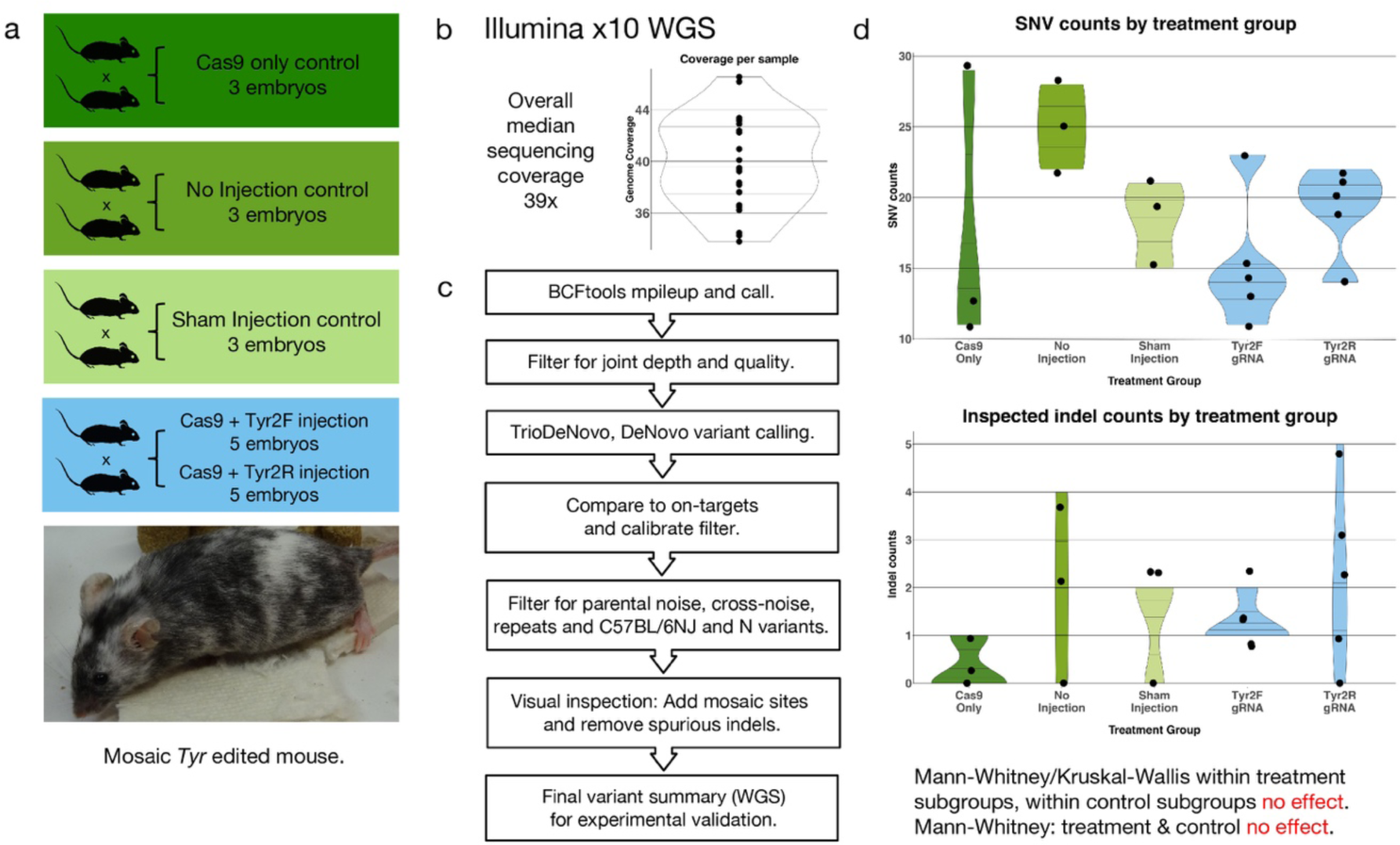
Analysis of CRISPR off-targets by whole genome sequencing. (a) Experimental design: Four sets of C57BL/6N parents gave rise to 9 control embryos (3 “no injection”, 3 ‘‘sham injection” with water only and 3 “Cas9 only”), and 10 treated embryos (5 were injected with Cas9 and Tyr2F gRNA and 5 were injected with Cas9 and Tyr2R gRNA).
(b) Whole genome sequencing: All 27 mice / embryos were subjected to whole genome sequencing with median depth 39.5x and an average of 3.4% of bases with read depth less than 11x.
(c) Variant calling and filtering: starting from the joint variant call (bcftools mpileup + bcftools call), a sequence of filter steps were performed to detect only *de novo* mutations and remove likely false positives arising from low-level parental mosaicism and repeat regions. Parental-noise: alternate-allele reads present in either parent. Cross-noise: alternate reads from all other (non-parental) samples.
(d) Filtered SNV and Indel counts are not significantly different within control groups, within treatment groups, or between control and treatment groups.

Sequencing was performed on the Illumina X10 WGS platform yielding a median sequencing depth of 39.5x per genome (**Fig. 1b**, Supplementary Table 2). In parallel, targeted Illumina MiSeq sequencing to a mean depth of 10,800 reads of the *Tyr* target site was performed to comprehensively profile mosaicism, with these data analyzed using the CRISPResso software[6] (Supplementary Table 3). MiSeq analysis revealed a targeting efficiency of over 85% for Tyr2R and mosaicism, with a median of two variants per embryo (Supplementary Table 3). Importantly, to ensure our experiment was representative of the many thousands of CRISPR experiments performed worldwide, including those of the International Mouse Phenotyping Consortium, we compared these data to MiSeq data from 324 mice mutagenized using the Tyr2R gRNA, revealing good concordance of targeting efficiency and mosaicism (Supplementary Table 3).

We next performed variant calling on the WGS data using bcftools mpileup and bcftools call[7], configured to be sensitive to low allele-fraction indels (insertions/deletions)(**Fig. 1c**). Starting with a median of 324,561 variants per sample, we filtered the results to ensure adequate depth (10x) and variant quality (10) in all samples, resulting in a median of 225,671 variants per sample. Candidate DNMs (those not inherited from either parent) in the embryos were called using the TrioDeNovo software[8], which resulted in a median of 6,852 variants per embryo (489 SNVs / 6,450 indels) (Supplementary Table 4). Prior to further filtration of these candidate DNMs, we first looked for the presence of mutations within 10bp of any candidate CRISPR off-target site for Tyr2F or Tyr2R, as defined by the Cas-OFFinder software[9]; allowing up to 3 mismatches with a 1 nucleotide DNA/RNA bulge, and up to 4 mismatches without a DNA/RNA bulge. Importantly, we found no such coincident sites in any embryos from the CRISPR-*untreated* group and only on-target variants in the CRISPR-treated group (Supplementary Table 5), suggesting that if there are recurrent CRISPR-induced off-target alterations they are exceedingly rare. We next applied a validated filtration strategy to refine our candidate DNM calls[10], removing false positives arising from mosaic alleles in either parent, as well as those in proximity to repeats. Alignments for all SNVs and indel variants were then inspected visually for the presence of mosaic alleles (i.e. a second alternative allele at the same locus). In the same way, all indels were visually inspected to remove further false positives. This resulted in a median of 19 SNVs and 1 indel per embryo (Table 1), which is broadly consistent with prior work that aimed to define the *de novo* mutation rates in mice[10]. All variants were validated with targeted MiSeq sequencing to a depth of at least 10,000 reads per locus (Supplementary Table 8).

**Table 1:** Initial and filtered *de novo* variant counts from whole genome sequencing. Summary table of initial variant counts, *de novo* variant counts, and filtered SNVs and Indels. Treatment Groups: Cas9 only (without gRNA), no injection (uninjected embryos), sham injection (water only), Tyr2R treated (Tyr2R gRNA + Cas9), Tyr2F treated (Tyr2F gRNA + Cas9). Variants passing basic depth / quality filters: bcftools joint call variant count per animal passing joint-depth and genotype quality filters (see Supplementary Methods). Candidate *de novo* mutations: all candidates produced by TrioDeNovo caller. Final SNVs / Final Indels: all SNVs/Indels remaining after filtering for false positives arising from low-level mosaicism, known C57BL/6NJ & C57BL/6N variants, proximity to UCSC repeat regions and further visual inspection.

| Mouse Sample | Treatment Group | Relationship | Variants passing Basic depth / Quality Filters | Candidate *de novo* Mutations | Final SNVs | Final Indels |
| --- | --- | --- | --- | --- | --- | --- |
| MD5617a | - | parent | 225,408 | - | - | - |
| MD5618a | - | parent | 229,105 | - | - | - |
| MD5630a | cas9 only | embryo | 226,692 | 7,106 | 13 | 0 |
| MD5631a | cas9 only | embryo | 216,975 | 6,533 | 11 | 0 |
| MD5632a | cas9 only | embryo | 217,421 | 6,626 | 29 | 1 |
| MD5619a | - | parent | 226,236 | - | - | - |
| MD5620a | - | parent | 230,156 | - | - | - |
| MD5624a | no injection | embryo | 228,022 | 7,377 | 25 | 4 |
| MD5625a | no injection | embryo | 219,537 | 6,599 | 22 | 2 |
| MD5626a | no injection | embryo | 217,827 | 6,510 | 28 | 0 |
| MD5616a | - | parent | 224,201 | - | - | - |
| MD5623a | - | parent | 221,461 | - | - | - |
| MD5627a | sham injection | embryo | 226,709 | 7,732 | 15 | 2 |
| MD5628a | sham injection | embryo | 225,793 | 7,693 | 21 | 2 |
| MD5629a | sham injection | embryo | 224,692 | 7,646 | 19 | 0 |
| MD5621a | - | parent | 228,855 | - | - | - |
| MD5622a | - | parent | 228,994 | - | - | - |
| MD5633a | Tyr2R treated | embryo | 228,026 | 7,083 | 19 | 0 |
| MD5634a | Tyr2R treated | embryo | 225,671 | 6,837 | 14 | 3 |
| MD5635a | Tyr2R treated | embryo | 231,888 | 7,308 | 22 | 5 |
| MD5636a | Tyr2R treated | embryo | 226,002 | 6,985 | 21 | 2 |
| MD5637a | Tyr2R treated | embryo | 221,699 | 6,618 | 20 | 1 |
| MD5638a | Tyr2F treated | embryo | 222,030 | 6,852 | 15 | 1 |
| MD5639a | Tyr2F treated | embryo | 219,136 | 6,440 | 13 | 2 |
| MD5640a | Tyr2F treated | embryo | 220,946 | 6,750 | 11 | 1 |
| MD5641a | Tyr2F treated | embryo | 222,174 | 6,803 | 14 | 1 |
| MD5642a | Tyr2F treated | embryo | 226,899 | 7,000 | 23 | 1 |
| **MEDIAN** |  |  | **225,671** | **6,852** | **19** | **1** |

A comparison of the expected variants at the *Tyr* locus detected by WGS and targeted MiSeq sequencing (Supplementary Table 6) shows that of the 20 indels detected by the MiSeq pipeline, 18 were also detected by the WGS pipeline, with the missing indels having low allele frequencies (7% and 7.5%, as defined by MiSeq sequencing). We are therefore confident that our WGS pipeline will detect genome-wide off-target damage with a range of allele fractions and mosaicism similar to on-target variants. Further, given our median depth (39.5x) and our minimum required *de novo* allele frequency (10%, Supplementary Methods), our power to detect a DNM occurring in the single-cell or two-cell stage of the zygote is at least 99%. Our power to detect a DNM with on-target variant allele fraction 0.17 (the median allele fraction seen in our experiments, Supplementary Table 6) is 85%.

Using the final counts of filtered SNVs and indels for each embryo (**Table 1**), we conducted a Kruskal-Wallis Rank test, detecting no significant difference in DNM counts between the “no injection”, “sham” and “cas9 only” untreated embryo groups (p=0.30 and p=0.37 for SNVs and indels, respectively). Similarly, a Wilcoxon Rank Sum test failed to detect significantly different SNV- or indel-DNM counts between the “Tyr2F” and “Tyr2R” CRISPR-treated groups (p=0.25 and p=0.43 for SNVs and indels, respectively). Based on these analyses (**Fig. 1d**), we combined variant calls from embryos in the two CRISPR-treated groups and in the same way combined data from the three untreated groups. Notably, using these data a Wilcoxon Rank Sum test failed to detect a significant difference in SNV or indel counts between the CRISPR-treated and untreated groups; p=0.30 and p=0.45, respectively (Supplementary Table 7).

Finally, we measured the impact of using unrelated parents on the false-positive DNM rate by deliberately choosing the parents of the Cas9-only embryos when analyzing all embryos in the study; the male parent (CBLT8902) was up to 7 generations removed from all other male parents (Supplementary Figure 1). Performing a comparable subset of filtrations and comparing variant counts by sample to the correctly analysed embryos at the same filtration point showed a median increase of 66 false variants per embryo (Supplementary Figure 1, Supplementary Table 9), highlighting the importance of using trios of mice when studying potential off-target rates.

## Conclusion

We conclude that if CRISPR mutagenesis were causing SNV or indel off-target mutations in treated embryos, then the number of these mutations is not statistically distinguishable from the background rate of DNMs occurring due to other processes. This work should support further efforts to develop CRISPR-Cas9 as a therapeutic tool.

## Acknowledgements

We thank William C. Skarnes and Allan Bradley for their scientific advice. All mouse work was undertaken by the Sanger Research Support Facility (RSF), with assistance from Evelyn Grau, Joanne Doran, Ellen Brown, Mike Woods and Catherine Tudor. Sequencing was performed by the Sanger DNA sequencing pipeline, with additional analysis by the Cancer Genome Project Informatics team. This work was supported by Wellcome funding.

## Author Contributions

KB, BD, ER performed the experiments. VI and MT performed the analysis. DJA oversaw the project. All authors contributed to the writing of the manuscript.

## Supplementary Methods

### 1. gRNA choices and characterisation of efficiency and mosaicism

We chose two gRNAs (Tyr2F and Tyr2R) within exon 2 of the Tyrosinase *(Tyr)* gene, as mutations at this locus do not cause lethality or influence embryogenesis. *Tyr* is responsible for black coat colour and eye pigmentation in wild type (WT) C57BL/6 (B6) mice and bi-allelic mutation of this gene results in complete loss of pigmentation (albinism)[4]. Therefore, detection of biallelic mutations as well as mosaicism caused by CRISPR/Cas9 activity is easily visible as a change in coat colour.

The Tyr2R gRNA was selected as previously described[11] and together with the Tyr2F gRNA represent a typical range of off-target scores. Off-target scores for the Tyr2F and Tyr2R gRNAs were determined for the default (NGG) PAM only, using the reference mouse genome (mm10) in the Cas-OFFinder tool[9]. Off-target scores were also determined for the Schaefer sgRNA#4[12] using the FVB/NJ mouse genome, to ensure that off-target scores were similar.

Summary of off-target scores.

| CRISPR | Sequence | Number of nucleotide mismatches |  |  |  |  | Off-target totals |
| --- | --- | --- | --- | --- | --- | --- | --- |
|  |  | 0 | 1 | 2 | 3 | 4 |  |
| Tyr2F | Target only | 1* | 0 | 0 | 4 | 51 | 55 |
|  | Target +1 (DNA bulge) | 0 | 3 | 5 | 75 | - | 83 |
|  | Target -1 (RNA bulge) | 2 | 2 | 8 | 245 | - | 257 |
|  |  |  |  |  |  |  | **395** |
| Tyr2R | Target only | 1* | 0 | 2 | 15 | 161 | 178 |
|  | Target +1 (DNA bulge) | 1 | 1 | 12 | 324 | - | 338 |
|  | Target -1 (RNA bulge) | 0 | 4 | 71 | 911 | - | 986 |
|  |  |  |  |  |  |  | **1502** |
| Schaefer sgRNA#4 | Target only | 1* | 0 | 0 | 10 | 129 | 139 |
|  | Target +1 (DNA bulge) | 0 | 2 | 8 | 226 | - | 236 |
|  | Target -1 (RNA bulge) | 1 | 3 | 55 | 923 |  | 982 |
|  |  |  |  |  |  |  | **1357** |

\* This indicates the *on-target* gRNA

The sequences for the gRNAs are as follows (PAM in bold):

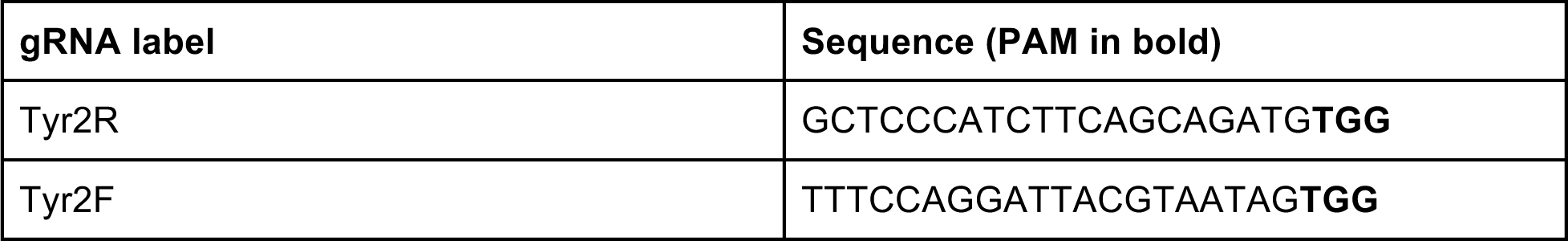

To determine the expected cutting efficiency and mosaicism from a large number of experiments we chose Tyr2R, comparing different Cas9 sources (mRNA or protein) as well as gRNA sources *(in vitro* transcribed or synthetic). The gRNA was mixed in RNase free water (Ambion) at a concentration of 25ng/ul together with either Cas9 mRNA or protein at 50ng/ul. The CRISPR reagents were injected into the cytoplasm of zygotes and F0 pups were scored for black, mosaic and albino coat colour. In addition, genomic DNA from earclips was extracted from F0 mice using the Sample-to-SNP kit lysis buffer (Life Technologies), sequenced using Illumina MiSeq and subsequently analysed with CRISPResso[6]. This enabled us to determine the targeting efficiency and how many different alleles were present within each founder animal, therefore making it possible to score mosaicism (more than one mutated allele detected in the animal) for each condition. The synthetic gRNA was the most efficient, with 70% of pups showing a mutant genotype, which were therefore used for the trio experiment (Supplementary Table 3).

For the trio experiments, synthetic gRNA consisting of crRNA and tracrRNA (Sigma) were diluted and mixed in RNase free water at equimolar ratios of 0.7pmol/ul each. Cas9 protein (obtained from Marko Hyvonen, Department of Biochemistry, University of Cambridge) was added to a working concentration of 50ng/ul and the mixture was incubated at 25^o^C for 10 minutes before zygote injection.

The following concentrations were injected for each of the experimental groups:

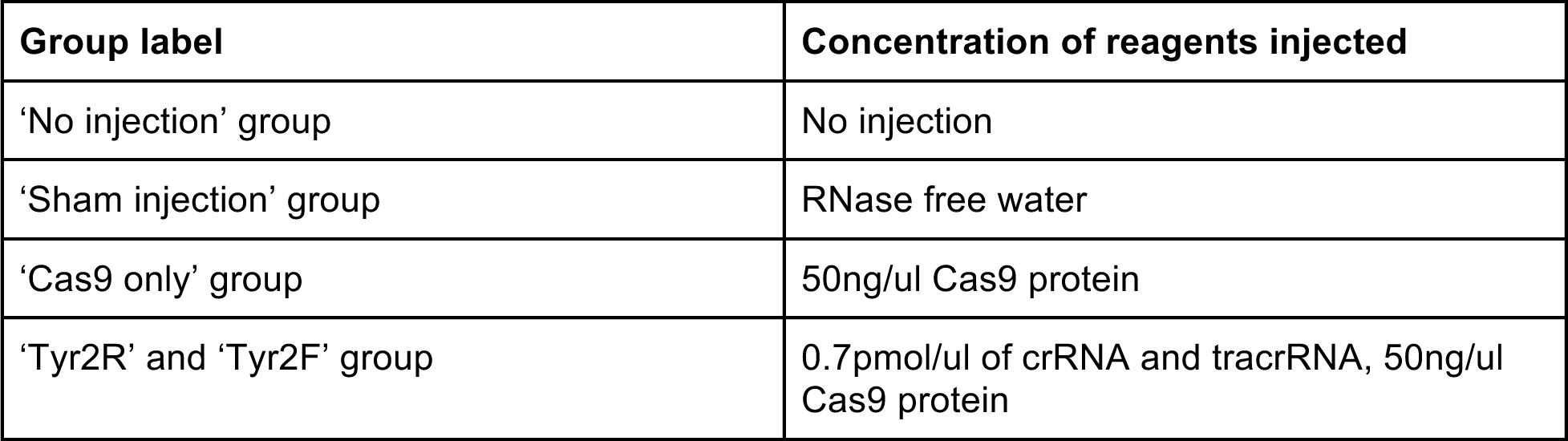

| **Group label** | **Concentration of reagents injected** |
| --- | --- |
| 'No injection' group | No injection |
| 'Sham injection' group | RNase free water |
| 'Cas9 only' group | 50ng/ul Cas9 protein |
| 'Tyr2R' and 'Tyr2F' group | 0.7pmol/ul of crRNA and tracrRNA, 50ng/ul Cas9 protein |

Genomic DNA was extracted from the kidney of the parents or 12.5 d.p.c. embryos using a DNeasy Blood & Tissue Kit (Qiagen) according to manufacturer’s instructions. DNA was quantified using a NanoDrop spectrophotometer, then used for sequencing using Illumina MiSeq and subsequently analysed with CRISPResso[6]. The results are presented in Supplementary Table 3 labelled with the treatment condition (e.g. ‘No injection’, etc), and are used to check the that the trio experiment is comparable to historical data at the target locus.

### 2. Zygote injection

4 ×4-week old C57BL/6NTac females were super-ovulated by intraperitoneal (IP) injection of 5 IU of pregnant mare’s serum (PMSG) at 11.00 hrs (on a 12hr light/dark cycle, on at 07:00/off at 19:00) followed 48hrs later by an IP injection of 5 IU human chorionic gonadotrophin (hCG) and mated overnight with C57BL/6NTac stud males. The next morning females were checked for the presence of a vaginal copulation plug as evidence of successful mating and females housed separately noting which males were used to plug each of the females. Oviducts were dissected one female at a time and harvests of cumulus masses kept separately in 4 different groups. Cumulus masses were released and treated with hyaluronidase as previously described[13]. Fertilized 1-cell zygotes were selected and maintained in KSOM media prior to cytoplasmic injection at 37^o^C in 4 separate dishes. MicroInjections were carried out between 23-25hrs post hCG. Cytoplasmic Injections were carried out as in Supplementary Table 1. The tyrosinase Tyr2F and Tyr2R microinjected embryo groups both came from the same zygote pool and hence had the same parentage. All other microinjection groups had a unique set of parents.

The Cas9 ribonucleoproteins (RNP) were backfilled into a microinjection needle. Microinjections were carried out using positive balancing pressure microinjecting into the cytoplasm of fertilized 1-cell zygotes held in FHM medium. A successful injection was indicated by visible movement in the cytoplasm after breaking the Oolemma. Microinjected 1 cell embryos were briefly cultured and viable zygotes were transferred the same day by oviducal embryo transfer into a 0.5 days post coital (d.p.c.) pseudo-pregnant female F1 (CBA/C57BL/6J) recipients[13]. After 12.5 d.p.c. recipient mice were humanely culled and embryos dissected and snap frozen. Previously, the parents from each group were humanely culled, tissue taken (liver, kidney and tail) and labelled according to which microinjection group they contributed to (Supplementary Table 1). All procedures performed in studies involving animals were in accordance with the ethical standards of the institution or practice at which the studies were conducted and performed with approval of the UK home office.

### 3. DNA extraction from mice and embryos

For the historical data, genomic DNA was extracted from earclips of F0 mice using the Sample-to-SNP kit lysis buffer (Life Technologies).

For the trio experiments, genomic DNA was extracted from the kidney of parent or 12.5 d.p.c. embryos using a DNeasy Blood & Tissue Kit (Qiagen) according to manufacturer’s instructions. DNA was quantified using a NanoDrop spectrophotometer.

### 4. Amplicon sequencing and analysis at the *Tyr* locus of mice and embryos

1ul of genomic DNA (from earclips of F0 founders, kidney of the parents or from 12.5 d.p.c. embryos) was used for amplicon specific PCR using genome specific primers (PE_tyrex2N_F1 and PE_tyrex2N_R1, see inset table), which flanked the expected on-target sites for Tyr2R and Tyr2F. The indexed libraries were sequenced using standard protocols and Illumina MiSeq technologies (Paired End 250bp runs).

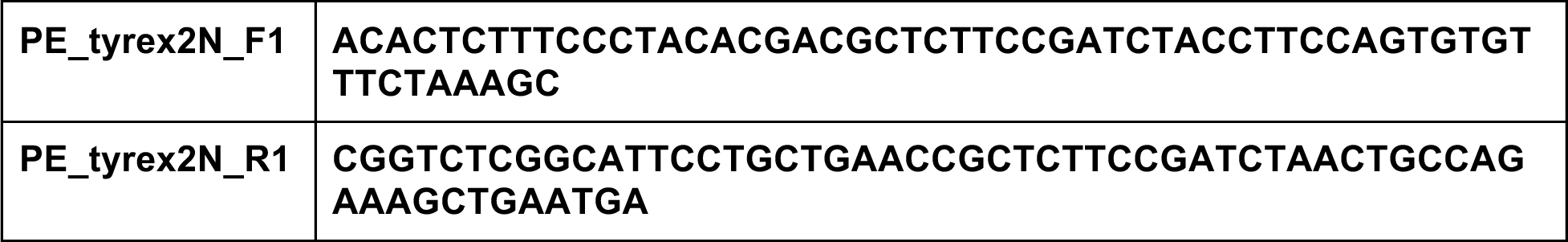

The paired-end fastq files generated were analysed using the CRISPResso sequence analysis package[6]. A typical command line used to run CRISPResso specified both gRNA sequences concurrently and allowed a window of 30bp around the gRNA sites to detect possible mutations: *CRISPResso -r1 sample_read1.fastq .fastq -r2 sample_read2.fastq -a <unedited_amplicon_sequence> -w 30 -g <Tyr2R_sequence>,<Tyr2F_sequence>.* The resulting sequence variants were visually inspected for indels with an allele fraction of at least 5%.

MiSeq analysis performed on the target region in these samples using CRISPResso showed that the parents, no injection, sham and Cas9 only groups showed no on-target activity, whereas the two gRNAs showed efficient cutting at their target site (Supplementary Table 3). Of the 10 mice showing targeting for the Tyr2R gRNA, 3 had one allele (30%), 5 had two alleles (50%) and 2 had three alleles (20%). Of the 7 mice showing targeting for the Tyr2F gRNA, 4 had two alleles (57%), with the remaining three mice containing either one, three or four alleles. This is in broad concordance with the historical data (Supplementary Table 3).

In order to cover a representative range of alleles at the on-target site we selected the following mice for WGS analysis: for the Tyr2R sample 1 mouse with one allele, 3 mice with two alleles and 1 mouse with three alleles; for the Tyr2F sample 1 mouse with one allele, 3 mice with two alleles and 1 mouse with three alleles.

### 5. Whole Genome Sequencing

Whole genome sequencing libraries were prepared using standard protocols for the Illumina X10 platform. The resulting sequence was aligned using ***bwa mem*** to the reference mouse GRCm38 assembly. The total mapped coverage varied from 34x to 47x, with a median of 39.5x (Fig. 1 b). The median fraction of bases with a coverage of greater than 50x was 39% and the median fraction of bases with a coverage of less than 11x was 3.4% (Supplementary Table 2).

### 6. Probability of detecting de novo mutations at low allele fractions (power)

To find our power to resolve mosaic DNMs, we note that a mosaic heterozygous DNM occurring in a two-cell zygote would yield 10 mutant reads from the (median) 39.5x read coverage. We model the actual number of mutant reads observed with a poisson distribution with parameter *lambda* = 10. Since our minimum required variant allele frequency is 10% (Supplementary Methods Section 7), we must observe at least 4 mutant allele reads out of 39 to call a DNM. The probability of seeing 3 or fewer reads in this case is given by the R function *ppois(3,10,lower.tail=TRUE)* = 0.01. This suggests that we should detect a DNM in a two-cell embryo approximately 99% of the time. By contrast, our median observed on-target variant allele fraction is 0.17 (median of column I, Supplementary Table 6). This allele fraction is represented by 6.6 variant reads out of 39.5 reads. The probability of seeing 3 or fewer reads in this case is given by *ppois(3,6,lower.tail=TRUE)* = 0.15. This suggests that we should detect a DNM in our samples approximately 85% of the time.

### 7. Variant calling, Trio Analysis Pipeline and Filtering of Candidate *De Novo* Mutations

Aligned sequence was jointly variant called for all parents and offspring using *bcftools mpileup, bcftools call, bcftools norm and bcftools filter[7].* The bcftools version and command options used are as follows: *bcftools-1.6 mpileup -a AD -C50 -pm2 -F0.05 -d10000, bcftools-1.6 call - vm, bcftools-1.6 norm -m -any, and bcftools-1.6 filter -m+ -sLowQual -e”%QUAL<=10” -g3 - G10.*

Note: a joint (multiallelic) variant call was performed on all parents and offspring at the same time, and that a sensitive bcftools mpileup setting (requiring only 2 or more indel reads and a minimum indel allelic fraction of 0.05) was used, in order to detect low-level indel mosaicism prior to any filtering. *bcftools* called between 317,235 and 332,048 variants per sample with a genotyping quality >=10, with a median of 324,561 variants (Supplementary Table 4). All variant loci were required to have a total depth of at least 10 reads for further analysis.

*De novo* mutation calling on all variants was performed by running TrioDeNovo software[8], using default settings, independently on each parent/offspring trio. TrioDeNovo produced between 6,440 and 7,732 candidate mutations per offspring, with a median of 6,852 mutations (Supplementary Table 4).

Based on the read depths and variant allele fractions seen in the expected on-target mutations called by TrioDeNovo, we filtered all candidate *de novo* variants with the following criteria, to remove likely false positives: (1) We required a minimum variant allele fraction of 10% to allow for mosaic alleles. (2) To avoid false positives arising from low-allele-fraction mosaic variants in either parent, we removed variants with *any* alternate-allele reads present in *either* (“parental noise”). (3) We removed any variant coincident with an allele reported in the C57BL/6NJ strain of the Mouse Genomes Project[14] or specifically in the C57BL/6N^*Tyr*^ strain^10^. (4) To avoid further false positives arising from low-allele-fraction mosaic variants present in the population, we allowed a maximum contribution of only 2% alternate reads from all other (non-parental) samples (“cross noise”). (5) As repeat regions can cause mis-alignment of reads resulting in false positive calls, we merged all individual UCSC repeat tracks and the UCSC Repeatmasker track (about 1.2Gb of sequence), and removed any variant inside or 1 bp adjacent to any merged repeat region. (6) We removed any *de novo* variant shared by two or more samples, as this would be extremely unlikely, and such mutations are more likely mosaic in the parents. Although it is possible this could remove a preferred off-target mutation, we note this removed only 3 variants altogether. (7) Every variant locus was inspected to check whether any position was actually mosaic (i.e. contained two or more alternative variants). These extra alternate variants were not consistently called and had to be manually re-inserted. (8) We found indel variants to be especially susceptible to false positive calls arising from un-annotated microsatellites or repeats, so we visually inspected all indel variants to remove any variant still in or adjacent microsatellites/homopolymers, which were not annotated by the UCSC repeat tracks. These filters resulted in 11 to 29 SNVs per sample (median 19) and 0 to 5 indels per sample (median 1). Every SNV and indel has been listed in Supplementary Table 8.

### 8. Intersection of all variants with potential off-target locations for Tyr2F and Tyr2R gRNAs

The CAS-OFFinder tool[9] was used to find all potential off-targets sites based on sequence homology to either the Tyr2F or Tyr2R gRNAs, allowing up to 3 mismatches with 1 bp of inserted or deleted sequence and up to 4 mismatches with no inserted or deleted sequence.

This resulted in 395 and 1,507 potential off-target sites being detected for Tyr2F and Tyr2R, respectively (Supplementary Table 5). These 20bp gRNA positions were intersected (allowing for a window size of 10bp) with *all* candidate *de novo* mutations before filtering, using *bedtools-2.23 window.* The results are presented in (Supplementary Table 5, worksheet “RGEN_DNM_10bp_overlap”).

### 9. Comparison of SNVs and Indel counts between treatment groups

Due to the small number of samples and the low variant counts in each group, we chose to perform either a Kruskal-Wallis Rank Sum test (in the case of more than two groups) or a Wilcoxon Rank Sum test (for only two groups), to assess the null hypothesis: namely that the counts in each group were drawn from the same population. Tests were performed using the R function *kruskal.test or wilcox.test.* The groups compared, the values tested, the test used and test results are presented in Supplementary Table 7.

### 10. Validation with PCR and sequencing

1ul of genomic DNA from 12.5 d.p.c. embryos or 0.5ul of genomic DNA from the kidney of the parents was used for amplicon specific PCR using genome specific primers (Supplementary Table 8). The indexed libraries were sequenced using standard protocols and Illumina MiSeq technologies (Paired End 250bp runs).

These SNVs and indels were then subject to validation by targeted MiSeq deep sequencing: each candidate position was sequenced to an average depth of at least 10,000 reads in embryos and (pooled) parent samples. The sequences were directly aligned to the GRCm38 assembly using *bwa-mem*, and the resulting alignments directly inspected at each variant location using *samtools-1.3.1 mpileup -d50000 -Q0 -q0* to check for the presence of the alternate allele in the parents’ sample, and to confirm the alternate allele in the embryo sample. Locations in the pooled parent sample with more than 100 reads showing the alternate allele and greater than 1% alternate allele fraction were classed as not validated, as were locations in the embryo samples showing less than 1% alternate allele fraction. We found that **87%** of SNVs and **73%** of indels were validated (Supplementary Table 8).

### 11. Re-analysis of De Novo Mutations with genetically distant parents

To estimate the effect of using distantly related parents to the measured *de novo* mutation rate, we re-analyzed our embryos for DNMs using incorrect parents. It can be seen from the relationships of matings contributing to our experiment’s parents and embryos (Supplementary Figure 1a) that the male parent of the “Cas9 only” embryos (mating CBLT8902) was distant by up to seven generations from the equivalent male parents of the other treatment groups (matings CBLT8762, CBLT9125, CLBT8712), whereas the female parents for all treatment groups were drawn from mating CBLT9125. We therefore chose to *fix* the parents for the “Cas9 only” group as the parents for *all* treatment groups as input to TrioDeNovo, and reran our pipeline, including filtration of variants up to the removal of C57BL6NJ/N repeats (filtration steps 1 - 4, section 6). It was not possible to run further filtration, e.g. removal of cross-animal noise, as the “correct” parents were present in the other animals.

The results show a median of 98.5 DNMs per animal for the treatment groups with distantly related controls (“No Injection”, “Sham injection”, “Tyr2F” and “Tyr2R”), whereas the treatment group with the correct control (“Cas9 Only”) has a median number of 28 DNMs per animal (Supplementary Figure 1, Supplementary Table 9). The equivalent median of all treatment groups is 32 (Supplementary Table 4, column L; ‘not on-target, vaf, parent noise, repeat and BL6NJ/N filtered’). This demonstrates the effect of using unrelated or distantly related parents as controls when searching for CRISPR off-targets, and reinforces the need for studies to use trios in these experiments.

## Supplementary Figure

**Supplementary Figure 1:**
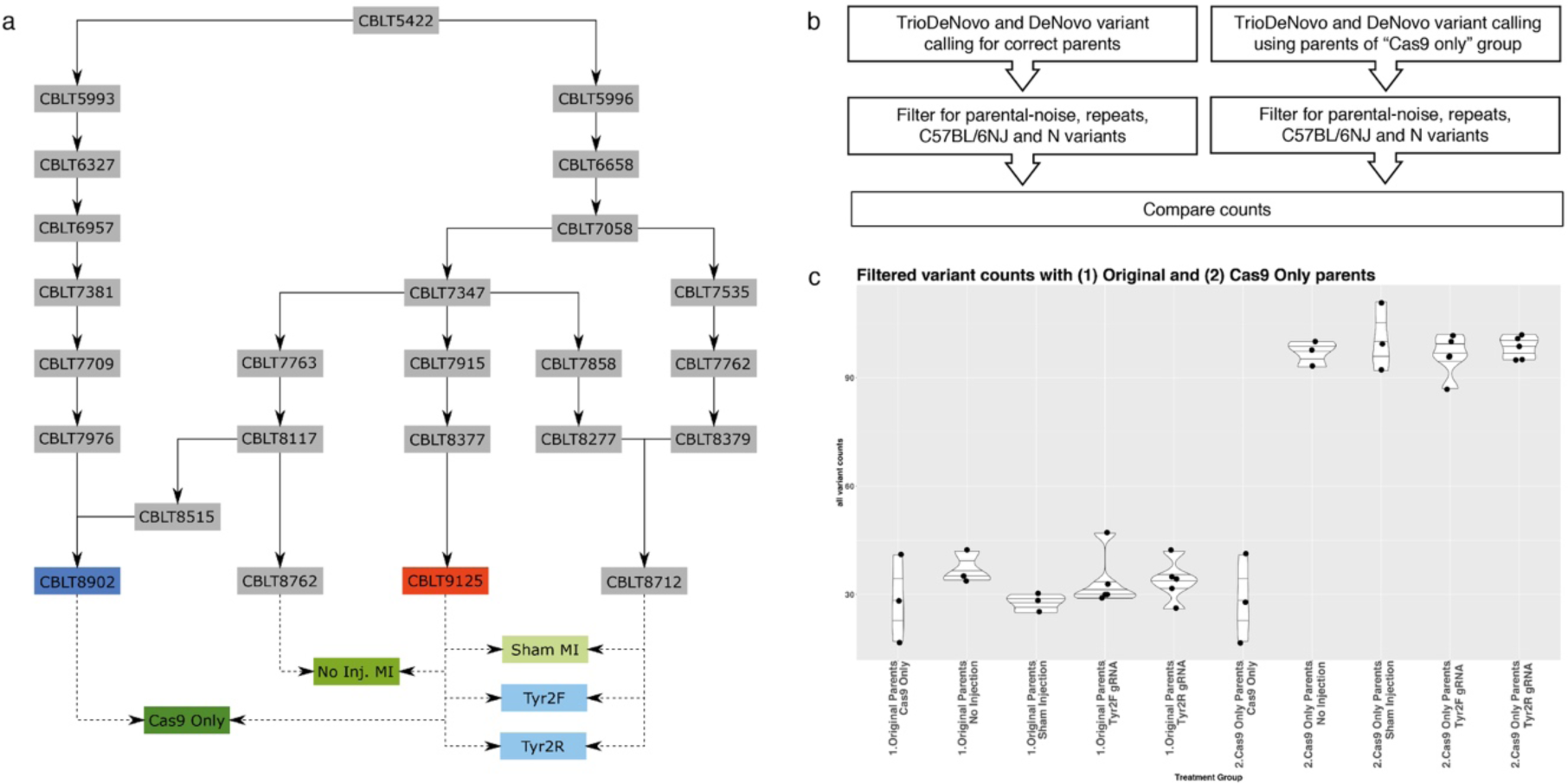
Analysis of parent effects on de *novo* mutations. (a) Network of heredity between the matings of mice from which mice parents were drawn. All female parents were drawn from mating CBLT9125 (red). Male parents were drawn from matings CBLT8712, CBLT8762 and CBLT8902 (dark blue) that is 7 generations distant. (b) Scheme of the filtration pipelines for both the “correct” calling approach and the “incorrect parent” calling approach, generating variant counts that can be compared. (c) Graphs of variants counts from “correct parent” pipeline and “incorrect parent” pipeline showing the effect of choosing distantly related parents on *de novo* variant calls: an average increase of 60 variants.

## Supplementary tables

Supplementary tables provided in excel format (see zip file).

**Supplementary Table 1: List of zygote injections**.

Counts of embryos microinjected, transferred and harvested at 12.5 d.p.c by treatment group. **Supplementary Table 2: Illumina X10 read coverage**.

**Worksheet: Total Depth**

Total numbers of bases sequenced by Illumina X10 and average depth by mouse sample.

**Worksheet: Depth by Coverage Bin**

For each sample the columns 1+, 11+ etc show the fraction of all sequenced bases at coverage greater >= 1x, >=11x, etc.

**Supplementary Table 3: Summary of mosaicism for determining cutting efficiency**.

Cutting efficiency and mosaicism obtained from Tyr2R, using different Cas9 sources (mRNA, protein) and different gRNA sources *(in vitro* transcribed or synthetic). Historical cutting efficiency is compared to the current experiment.

**Worksheet: Efficiency and Mosacism**

Shows cutting efficiency and mosaicism rates from both historical mouse data (Rows “Cas9 Protein” and “Tyr2R RNP”) and the present experiment (Rows “parent”, “No Injection”, “Sham injection”, “Cas9 only”, “Tyr2F”, “Tyr2R” describe each treatment condition). Historical data shows broad similarity to current data conditions “Tyr2R” and “Tyr2F” for targeting efficiency (column E) and average variants per embryo (column G).

**Columns**:

Targeting efficiency: the number of animals displaying a mutant genotype with allele frequency > 5%. Percentage is stated of the number of pups analysed.

Number with Mosaic genotype: The number of animals or embryos displaying multiple alleles at the same target site. Percentage is stated of the number of pups analysed.

Average variants per embryo: the average number of distinct alleles per embryo or mouse in each historical or current treatment group.

**Worksheet: TestsOfDifferentConditions**

Historical data. Shows the expected cutting efficiency and mosaicism of Tyr2R under different Cas9 sources and gRNA sources. Synthetic gRNA was most efficient with 57 out of 82 pups born showing indels (70%). On average, 53% of mice showed one allele, 33% two alleles, 12% three alleles, 2% four alleles and 0.5% five alleles. As the synthetic gRNA showed the highest cutting efficiency, we used these for further experiments.

**Supplementary Table 4: Total variant counts and filtered counts by sample**.

This is a detailed version of Table 1, with further details on the reduction of variant counts as successive filters are applied. (see Supplementary Methods). Samples are grouped into parent / offspring groupings.

**Columns:**

Sample: sample label of each parent or embryo

Treatment Group:

Cas9 only - gRNA omitted in injection

No injection - uninjected embryos

Sham injection - injected with water

Tyr2R treated - treated with Tyr2R gRNA + Cas9

Tyr2F treated- treated with Tyr2F gRNA + Cas9

Quality passed: Variants per sample passing basic genotype quality > 10

Depth and quality passed: variant count per animal passing joint-depth (>10) and genotype quality filters

TrioDeNovo, depth, quality: all candidates produced by TrioDeNovo caller passing the previous two thresholds.

TrioDeNovo, depth, quality SNVs / Indels: breakdown of TrioDeNovo column by variant class.

On target: Indel variants produced by TrioDeNovo lying in the expected Tyr mutation regions

Not on-target, vaf and parent noise filtered: all SNVs/Indels *not* in the Tyr locus, remaining after filtering for false positives arising from parental mosaicism and minimum Variant Allele Fraction (>=0.1)

Not on-target, vaf, parent noise and repeat filtered = Variants passing prior filters which are not in or 1bp adjacent a UCSC repeat region.

Not on-target, vaf, parent noise, repeat and BL6NJ/N filtered = Variants passing prior filters which are not known BL6NJ or BL6N variants.

Not on-target, vaf, parent /cross noise, repeat and BL6NJ/N filtered: Variants passing prior filters which are not present at more than 2% in any other samples.

… Without shared DNVs: Variants passing prior filters, which are not shared between any two embryos:

Final SNVs: SNV variants passing all previous filters (and manual re-addition of any mosaic variant at same locus, if it exists).

Filtered Indels: Indel variants passing all previous filters (and manual re-addition of any mosaic variant at same locus, if it exists).

Final indels: indel variants passing all previous filters, visually inspected for correctness.

**Supplementary Table 5: Off-target locations with adjacent *de novo* mutations.**

**Worksheets: RGEN_Tyr2F_Offtarget_Sites, RGEN_Tyr2R_Offtarget_Sites: All expected CRISPR off target locations for Tyr2F and Tyr2R**.

Columns state Chromosome, Position and Direction of expected off-target site, as well as the extent of the mismatch (number of basepair mismatch and whether there is a “bulge” - i.e. a 1 bp insertion or deletion, and whether the “bulge” is a DNA or RNA bulge).

**Worksheet: RGEN_DNM_10bp_overlap_by_sample**

The intersection (with a 10bp window) of unfiltered *de novo* variant calls with all Tyr2F and Tyr2R gRNA candidate off-target locations. This shows that no untreated animals have any overlap within 10bp of a candidate Tyr2R or Tyr2F off-target site, and that only the on-target indels overlap the expected on-target site in the treated animals.

**Supplementary Table 6: WGS and MiSeq on-target alleles for comparison**.

Shows a comparison of on-target variants revealed by high-depth MiSeq sequencing and CRISPResso analysis (cutoff allele frequency 5%), compared to the variants at the same locations revealed by whole-genome X10 sequencing and bcftools analysis. We note that CRISPResso found no on-target variants in the untreated samples (MD5624a-MD5632a), so these samples are not presented.

CRISPResso found 20 variants in the Tyr-treated variants (MD5633a-MD5642a), with variant allele fractions ranging from 0.1 to 0.33, and with a typical mosaicism of 2 alleles per embryo. This confirms that the *Tyr* gRNAs are active in the treated samples, and that the activity at the on-target locations measured by mosaicism and allele fraction are within expected ranges. Deep-sequencing of the on-target location by MiSeq therefore shows mosaic indels in treated animals, and no mutation in controls.

X10 sequencing confirms both location and mosaicism in treated animals, with the exception of three indels, two of which are at low allele fraction (7% and 7.5%), and the third which is a second mosaic allele at *exactly* the same location as a called allele. Our pipeline manually adds in such mosaic alleles.

**Supplementary Table 7:Results for KrusKal-Wallis and Wilcoxon Rank Sum tests**

There are three control groups, containing three samples each: a “no injection” group, a “sham” injection group (injected with water) and a “Cas9-only” group. Due to the small number of samples in each group we elected to perform a Kruskal-Wallis Rank Sum test to detect any differences between these three groups. This test failed to detect any difference in SNV counts between groups - i.e. to reject the null hypothesis (chi-squared = 2.4, df = 2, p-value = 0.3012) and failed to detect any difference in indel counts between groups (chi-squared = 1.9874, df = 2, p-value = 0.3702).

The Tyrosinase-treated groups were either Tyr2R-treated (5 samples) or Tyr2F-treated (5 samples). We performed a Wilcoxon Rank Sum test and again failed to detect any difference between the groups for SNV counts (W = 6.5, p-value = 0.2492) or indel counts (W = 8.5, p-value = 0.4338).

Finally, based on these results, we merged all three untreated groups together and both *Tyr*- treated groups together. Using the Wilcoxon Rank Sum test, we were unable to detect any difference in SNV counts between the Tyr-untreated and Tyr-treated groups (W = 58, p-value = 0.3059) or indel counts (W = 35.5, p-value = 0.4471).

**Supplementary Table 8: Filtered variant positions for experimental validation**.

This lists the genomic location of all final DNM variants (SNVs and Indels) passing all filters and subsequently sent for validation, as well as PCR primer pairs used for validation, and the validation test results.

**Supplementary Table 9: Filtered variant counts for alternative *de novo* variant analysis**.

Lists filtered variant counts for an alternative *de novo* variant analysis in which the parents of the Cas9-only trio were deliberately set to be the parents for *all* embryos. *De novo* variant calling and filtration was carried out as before, with filtration stage stopped after the removal of parental mosaicism, proximity to repeats and the overlap with known BL6NJ/N variants. (This is the last sensible filter stage suitable for a comparison of this approach with the original data.) Column G lists the combined SNV and indel counts at this filtration point. As expected, the “Cas9 only” group - which was analysed using the correct parents - has significantly lower values of variants than the other groups. Column G is directly comparable with Supplementary Table 4 Column L (“not on-target, vaf, parent noise, repeat and BL6NJ/N filtered”).

